# Reelin engages non-canonical signaling pathways to drive endothelial remodeling and plasticity

**DOI:** 10.64898/2026.04.08.717341

**Authors:** Dante Maria Stea, Sofia Nutarelli, Maria Teresa Viscomi, Luca Madaro, Antonio Filippini, Alessio D’Alessio

**Affiliations:** Dipartimento di Scienze della Vita e Sanità Pubblica, Sezione di Istologia ed Embriologia; Università Cattolica del Sacro Cuore, Rome, Italy; Fondazione Policlinico Universitario “Agostino Gemelli”, IRCCS, 00168 Roma, Italy; Dipartimento di Scienze Anatomiche, Istologiche, Medico Legali e dell’Apparato Locomotore, Sezione di Istologia ed Embriologia Medica; Sapienza Università di Roma, Rome, Italy

## Abstract

**BACKGROUND:** The vascular endothelium is a dynamic tissue central to vascular homeostasis and disease, with endothelial cells (ECs) exhibiting plasticity that drives adaptive remodeling. Reelin, a secreted extracellular matrix glycoprotein critical for neuronal migration via ApoER2/VLDLR-DAB1 signaling, may also modulate vascular function and inflammation. However, its direct role in EC biology remains unclear. We investigated Reelin as a context-dependent signaling modulator in ECs, assessing its engagement of non-canonical pathways and regulation of endothelial plasticity relevant to cardiovascular pathology.

**METHODS:** Human endothelial cells were stimulated with recombinant Reelin and analyzed by immunoblotting, immunofluorescence, and functional assays. Time-course studies assessed signaling, including phosphorylation of FAK, AKT, and DAB1 by Western blotting, while wound-healing assays quantified endothelial migratory capacity in vitro systems.

**RESULTS:** Reelin rapidly robustly activated noncanonical signaling in endothelial cells, increasing FAK and AKT phosphorylation in a time-dependent manner consistent with cytoskeletal remodeling. Canonical DAB1 activation was limited. Functionally, Reelin enhanced migration, upregulated Endoglin/CD105, and induced a remodeling-associated phenotype. Reelin silencing altered endothelial phenotype, clearly indicating a role in homeostasis. Signaling was independent of VEGFR2 interaction. Overall, Reelin preferentially engages FAK/AKT pathways to drive partial phenotypic modulation without full endothelial-to-mesenchymal transition.

**CONCLUSION:** We show that Reelin is a previously unrecognized regulator of endothelial signaling and plasticity, acting via non-canonical FAK- and AKT-dependent pathways. By partially and dynamically modulating endothelial phenotype, Reelin promotes a remodeling-permissive state without triggering full mesenchymal transition. These findings identify Reelin as a novel modulator of endothelial function with potential implications for vascular remodeling and cardiovascular disease.

**What Are the Clinical Implications?:** Our findings identify Reelin as a modulator of endothelial signaling with a clear bias toward non-canonical FAK- and AKT-dependent pathways that regulate endothelial plasticity and remodeling. This signaling profile is highly relevant to vascular diseases in which endothelial dysfunction is driven by maladaptive cytoskeletal reorganization, altered migration, and persistent activation rather than complete loss of endothelial identity. The ability of Reelin to promote partial and dynamically regulated phenotypic modulation suggests that it may operate at early and potentially reversible stages of vascular pathology. In this context, dysregulated Reelin signaling could contribute to pathological vascular remodeling, including processes underlying atherosclerosis, fibrosis, and microvascular dysfunction. These results also raise the possibility that circulating or locally produced Reelin may serve as an indicator of endothelial activation state, providing a novel biomarker for vascular disease progression. Importantly, the identification of a signaling bias toward FAK- and AKT-dependent pathways highlights potential therapeutic targets downstream of Reelin that could be selectively modulated to limit maladaptive endothelial remodeling while preserving essential endothelial functions. Collectively, this study positions Reelin signaling as a previously unrecognized and potentially actionable pathway in the regulation of endothelial behavior, with direct implications for the development of targeted strategies aimed at preventing or attenuating cardiovascular disease progression

## INTRODUCTION

The vascular endothelium is formed by a continuous monolayer of squamous ECs lining the luminal surface of all blood and lymphatic vessels ^1-4^. As the fundamental structural and functional unit of the vascular wall, ECs play a central role in maintaining cardiovascular homeostasis ^5^. Their remarkable phenotypic plasticity enables the dynamic plasticity of vascular permeability, hemodynamics, immune surveillance, and hemostasis, allowing rapid adaptation to physiological and pathological stimuli. Through the production of vasoactive mediators such as nitric oxide and endothelin-1 ^6-8^, ECs finely regulate vascular tone and blood flow, while their barrier function controls the selective exchange of nutrients, metabolites, and signaling molecules between the circulation and surrounding tissues. In addition, ECs actively coordinate immune responses by regulating leukocyte adhesion and transmigration and by secreting cytokines and chemokines that influence inflammation, angiogenesis, and tissue repair ^2, 9^. ECs also preserve thromboresistance by balancing coagulation, platelet adhesion, and fibrinolysis ^10, 11^. Endothelial dysfunction represents an early and pivotal event in the pathogenesis of cardiovascular disease. Characterized by reduced nitric oxide bioavailability, oxidative stress, inflammatory activation, and impaired barrier integrity, endothelial dysfunction disrupts vascular homeostasis and promotes a pro-atherogenic, pro-thrombotic state ^8, 12, 13^. Extensive experimental and clinical evidence identifies endothelial dysfunction as a reversible determinant of cardiovascular risk ^14, 15^, positioning the endothelium as both a mechanistic hub of vascular pathology and a critical therapeutic target ^16, 17^. Among the mechanisms contributing to sustained endothelial dysfunction, endothelial-to-mesenchymal transition (EndMT) has emerged as an important driver of maladaptive vascular remodeling ^18-20^. EndMT encompasses a spectrum of phenotypic changes in which ECs progressively lose endothelial identity and junctional organization while acquiring mesenchymal traits, including enhanced migratory capacity, contractility, and extracellular matrix production. Increasing evidence implicates EndMT and partial EndMT-like phenotypic shifts in atherosclerosis, vascular fibrosis, and microvascular disease, linking chronic inflammatory and metabolic stress to persistent erosion of endothelial phenotype. Reelin is a large secreted extracellular matrix glycoprotein best known for its essential role in neuronal migration and cortical lamination during development ^21, 22^. Canonically, Reelin signals through lipoprotein receptors Apolipoprotein E receptor 2 (ApoER2) and Very-low-density lipoprotein receptor (VLDLR), leading to phosphorylation of the adaptor protein Disabled-1 (DAB1) and activation of downstream pathways regulating cytoskeletal organization, cell survival, and junctional stability ^23-25^. In addition to this classical signaling axis, Reelin can activate non-canonical, DAB1-independent pathways ^26, 27^ involving integrins, FAK kinase, Src family kinases, and Rho GTPases, thereby modulating cell adhesion and migration ^23, 28^. While dysregulation of Reelin signaling has been extensively linked to neurological disorders, ^29-32^, its role in vascular endothelial biology remains poorly defined. Emerging evidence suggests that circulating Reelin may influence endothelial function and vascular remodeling, raising the possibility that Reelin signaling contributes to cardiovascular pathology ^33-35^. However, whether Reelin directly modulates endothelial plasticity or promotes EndMT-associated phenotypic changes has not been investigated. Here, we demonstrate that Reelin activates non-canonical signaling in human ECs, characterized by FAK and AKT phosphorylation, enhanced migration, and modulation of EndMT-associated markers. Our findings indicate that Reelin promotes an increased state of endothelial plasticity with features consistent with partial EndMT-like modulation rather than complete mesenchymal transition. To our knowledge, this is the first study identifying Reelin as a modulator of endothelial plasticity and EndMT-like features. These findings uncover a previously unrecognized extracellular signaling mechanism with potential relevance for vascular remodeling and the progression of cardiovascular disease.

## Methods

### Cell cultures and chemicals

Human-derived EA.hy926 ECs ^36^ were maintained in Dulbecco’s modified Eagle’s medium (Life Technologies Corporation) containing 10% (v/v) fetal calf serum, 2% (w/v) HAT (hypoxanthine/aminopterin/thymidine) (Sigma), 200µM L-glutamine, 100 units/ml penicillin/streptomycin (Life Technologies) at 37°C in a 5% CO2 humidified atmosphere. Human umbilical vein ECs (HUVECs) from a single donor obtained from Lonza (Walkersville, MD, USA) were maintained in endothelial basal medium (EBM) supplemented with EGM-2 media kit (Lonza).Upon reaching about 80% confluence, cells were detached using 0.05% trypsin and 0.02% EDTA solution at 37°C, centrifuged at 1200 RPM for 8 minutes and then replated with fresh complete culture medium. HUVEC within the initial 7-8 passages were utilized for subsequent analyses. VLDLR and ApoER2/LRP8 antibodies were from OriGene Technologies, Inc. (Rockville, MD, USA), VEGFR2, Phospho-FAK, CD105/Endoglin, and VE-cadherin antibodies were from Cell Signaling Technology (Danvers, MA, USA). Horseradish peroxidase (HRP)-conjugated monoclonal mouse anti β-Actin was from Sigma-Aldrich Inc. (St. Louis, MO, USA). HRP-conjugated secondary antibodies were from Bio-Rad Laboratories (Hercules, CA, USA). Oligofectamine Transfection Reagent was purchased from Thermo Fisher Scientific (Waltham, MA USA).

### Small Interfering RNA Experiments

Predesigned Dicer-substrate interfering RNAs (DsiRNAs) targeting the coding sequence of human Reelin cDNA (NM_005045) were synthesized and obtained from Integrated DNA Technologies (Coralville, IA, USA). To investigate the role of endogenous Reelin under physiologically relevant conditions, gene silencing was performed by transfecting HUVECs with 60 nM siRNA using Oligofectamine Transfection Reagent (Thermo Fisher Scientific), in accordance with the manufacturer’s protocol, in a total volume of 1 mL. To account for non-specific effects of siRNA delivery, cells were transfected with a commercially available non-targeting siRNA (Integrated DNA Technologies). At 5 hours post-transfection, 2 mL of fresh complete endothelial growth medium (EGM) were added, and cells were cultured for an additional 72 hours prior to downstream treatments and analyses. A commercially available non-targeting siRNA (Integrated DNA Technologies) was used as a negative control.

### Quantitative real time PCR analysis

For real time PCR analysis total RNA was extracted with the TRIZOL Reagent (Life Technologies Corporation), according to the manufacturer’s instructions. In brief, 3 µg of RNA were retro-transcribed into single-stranded DNA by a standard 20µl RT reaction with the TaqMan RT Reagent (Thermo Fisher Scientific Inc.). cDNA generated from the reverse transcription reactions was amplified by PCR with the SensiFAST SYBR Hi-ROX Kit from Meridian Life Science (Memphis, Tennessee, USA) in a total volume of 25 µl according to the manufacturer’s instructions. The level of gene was expressed as relative fold change vs. either the β-actin messenger RNA or Hypoxanthine Guanine Phosphoribosyltransferase (HPRT) using the ΔΔCt method using the Step One System Software (Thermo Fisher Scientific). Analysis of gene expression was performed with commercially available PrimeTime Predesigned qPCR Assays (Integrated DNA Technologies) for human VLDLR, ApoER2/LRP8, β-Actin and HPRT.

### SDS-PAGE, western blot, and immunoprecipitation analysis

Following treatments, ECs underwent two washes in ice-cold phosphate-buffered saline (PBS) and were then incubated in 1X Cell Lysis Buffer (Cell Signaling Technology) containing 20 mM Tris-HCl (pH 7.5), 150 mM NaCl, 1 mM Na2EDTA, 1 mM EGTA, 1% Triton, 2.5 mM sodium pyrophosphate, 1 mM beta-glycerophosphate, 1 mM Na3VO4, 1 µg/ml leupeptin, 2% SDS, and 1mM protease and phosphatase inhibitor cocktail for 15 minutes on ice. The whole cell lysate was clarified by centrifugation at 15,000 g for 15 minutes at 4°C, and the resulting supernatant was collected for further analysis. Protein content was quantified using the BCA Protein Assay Reagent (Thermo Fisher Scientific Inc.) and equal amounts of proteins were separated via SDS-PAGE, transferred onto 0,45-micron nitrocellulose membrane (Thermo Fisher Scientific Inc.), and subjected to immunoblotting. Detection of the primary antibody binding was achieved using Clarity Western ECL Substrate (Bio-Rad). Immunoblotted band intensities were measured using the ChemiDoc Imaging System (Bio-Rad) and analyzed with Image Lab software (Bio-Rad). The intensity of specific bands was normalized to that of the β-actin.

### Wound healing assay

EC migratory capacity was assessed using a monolayer scratch (wound-healing) assay ^37^. Confluent EAhy.926 monolayers cultured in 35-mm dishes were mechanically wounded with a P200 pipette tip to generate linear, uniformly sized cell-free gaps. The wounded monolayer was initially imaged by phase contrast microscopy at time 0. After gentle washing with PBS to remove detached cells and debris, cultures were incubated in fresh medium in the absence or presence of recombinant human Reelin at 500 ng/mL for about 18 hours. Cells were gently washed in PBS and wound closure was monitored by phase-contrast imaging using an inverted microscope equipped with a digital camera.

### Statistics

All quantitative data are expressed as the mean ± standard deviation (SD) from independent biological replicates. Densitometric analyses of immunoblots were performed using Image Lab Software (Bio-Rad Laboratories, Inc., Hercules, CA, USA), with target protein expression normalized to the corresponding loading control (β-actin) and expressed relative to untreated controls. Statistical comparisons among multiple groups were performed using one-way analysis of variance (ANOVA). Normality of data distribution was assessed prior to parametric testing. A p-value < 0.05 was considered statistically significant. Statistical analyses were performed using Microsoft Excel (Microsoft Corp., Redmond, WA, USA), and graphs were generated using SigmaPlot (Systat Software, Inc., San Jose, CA, USA).

## RESULTS

### The Reelin’s signaling is functional in human ECs

To explore the potential involvement of the Reelin pathway in human ECs, we first assessed the expression of the principal Reelin receptors, VLDLR and ApoER2, in cultured ECs. Western blot analysis followed by densitometric analysis demonstrated that both receptors were readily detectable under basal conditions (Figure 1A), with no statistically significant difference between VLDLR and ApoER2 expression levels (Figure 1B), indicating that ECs express comparable amounts of both receptors. These findings demonstrate that ECs possess the molecular machinery required to respond to Reelin, supporting a potential role for this pathway in endothelial biology relevant to vascular homeostasis and disease. In addition, incremental stimulation with recombinant Reelin did not significantly alter total ApoER2/LRP8 or VLDLR protein levels across the concentrations tested (Figure 1C), indicating that receptor abundance is not regulated at the total protein level in response to short-term Reelin exposure in ECs. These data suggest that receptor availability is maintained under the experimental conditions employed.

**Figure 1.**
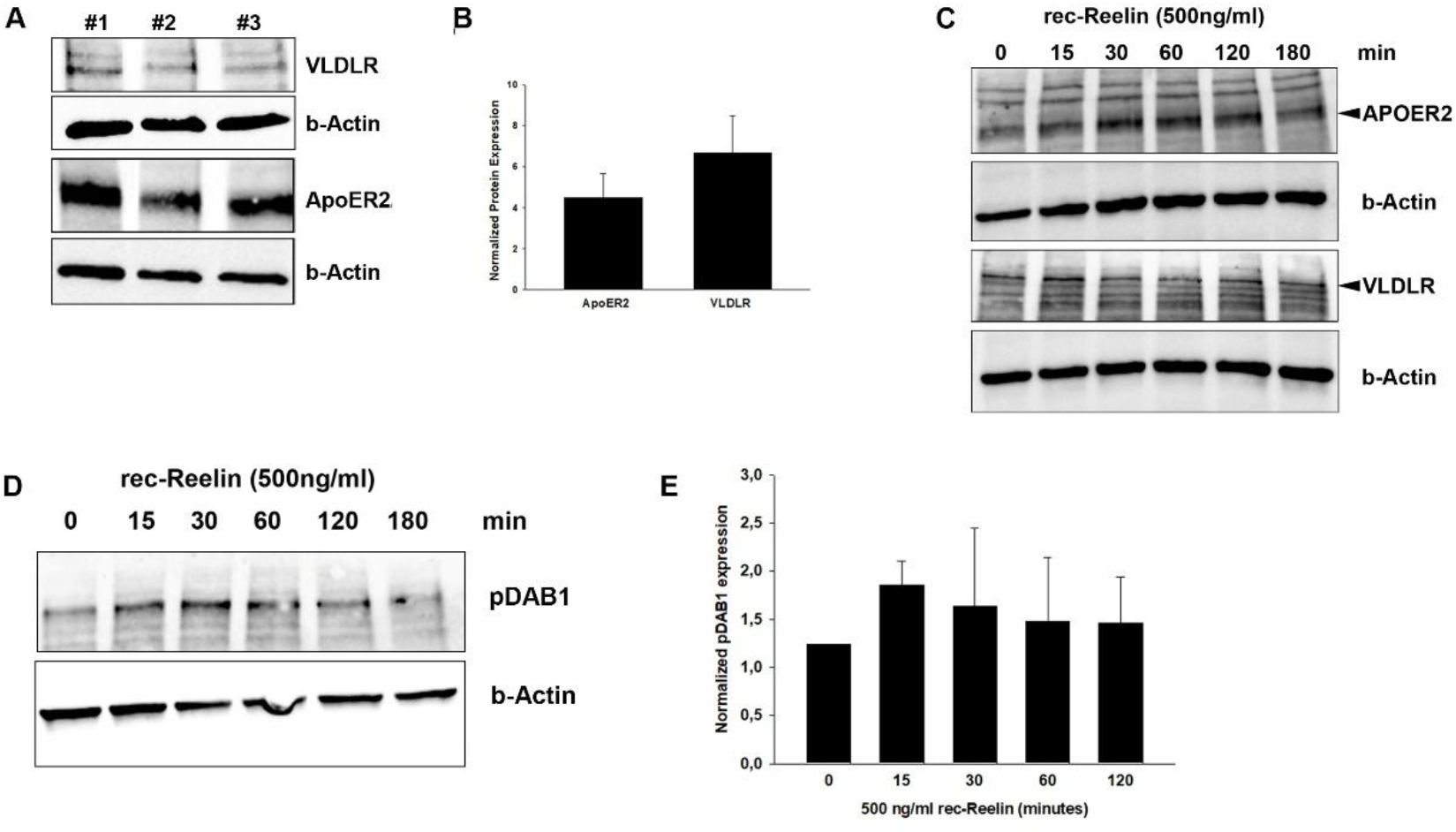
Expression of Reelin receptors and activation of DAB1 signaling in human ECs. (A) Representative immunoblots showing basal expression of VLDLR and ApoER2 in cultured human ECs (three independent replicate, #1–#3). β-Actin was used as loading control. (B) Densitometric quantification of basal ApoER2 and VLDLR protein levels normalized to β-actin. (C) Representative immunoblots showing total ApoER2 and VLDLR protein levels in ECs stimulated with recombinant Reelin (500 ng/mL) for the indicated time points (0–180 min). β-Actin served as loading control. (D) Representative immunoblot of phosphorylated DAB1 (pDAB1) in ECs treated with recombinant Reelin (500 ng/mL) for the indicated times. β-Actin was used as loading control. (E) Densitometric quantification of pDAB1 levels normalized to β-actin and expressed relative to control (0 min), (one-way ANOVA, p < 0.01), with a peak response observed at 30 minutes (p = 0.0035 vs. control). Data are presented as mean ± SD from independent experiments.

To determine whether Reelin activates canonical downstream signaling in ECs, we examined phosphorylation of DAB1, the central adaptor protein of classical Reelin signaling. Quantitative densitometric analysis revealed a statistically significant increase in DAB1 phosphorylation following stimulation with 500 ng/mL recombinant Reelin. The response was detectable as early as 15 minutes and peaked at 30 minutes compared to control (one-way ANOVA; p = 0.0035), followed by a sustained but modest elevation up to 180 minutes (Figure 1C). Although the magnitude of DAB1 activation was moderate compared with canonical neuronal systems, the kinetics were consistent with receptor-driven signaling. These findings indicate that Reelin is capable of engaging DAB1 phosphorylation in ECs, albeit with limited amplitude, suggesting partial activation of the canonical pathway under the experimental conditions employed.

### Reelin preferentially activates FAK- and AKT-dependent signaling pathways in ECs relevant to vascular remodeling

Although recombinant Reelin induced a modest but statistically significant increase in DAB1 phosphorylation, the magnitude of this response was limited compared with the robust activation of downstream signaling pathways implicated in endothelial remodeling. We therefore examined whether Reelin preferentially engages non-canonical signaling mechanisms associated with cytoskeletal dynamics and vascular adaptation. Stimulation of ECs with recombinant Reelin (500 ng/mL) elicited a rapid and sustained increase in AKT phosphorylation (Figure 2A). Phospho-AKT levels were detectable at early time points and remained elevated throughout the duration of stimulation, consistent with activation of pro-survival and pro-migratory signaling pathways known to regulate endothelial adaptation during vascular remodeling and cardiovascular disease. Concomitantly, Reelin markedly enhanced phosphorylation of full-length FAK (∼125 kDa), a central mediator of integrin-dependent signaling and endothelial mechanotransduction (Figure 2B). The increase in phospho-FAK was time-dependent and reproducibly observed following Reelin exposure, supporting activation of focal adhesion signaling pathways that govern cytoskeletal organization, adhesion turnover, and migratory responses. Importantly, Reelin treatment did not induce caspase activation (data not shown), arguing against apoptotic signaling ^38^ and reinforcing the interpretation that FAK activation reflects dynamic remodeling rather than stress-induced proteolysis. Collectively, these findings indicate that, although Reelin can engage elements of the canonical pathway, signaling output in ECs is strongly biased toward FAK- and AKT-dependent mechanisms. The preferential activation of pathways governing focal adhesion dynamics and cytoskeletal remodeling highlights a non-neuronal signaling architecture through which Reelin may contribute to endothelial plasticity and vascular remodeling processes relevant to cardiovascular disease.

**Figure 2.**
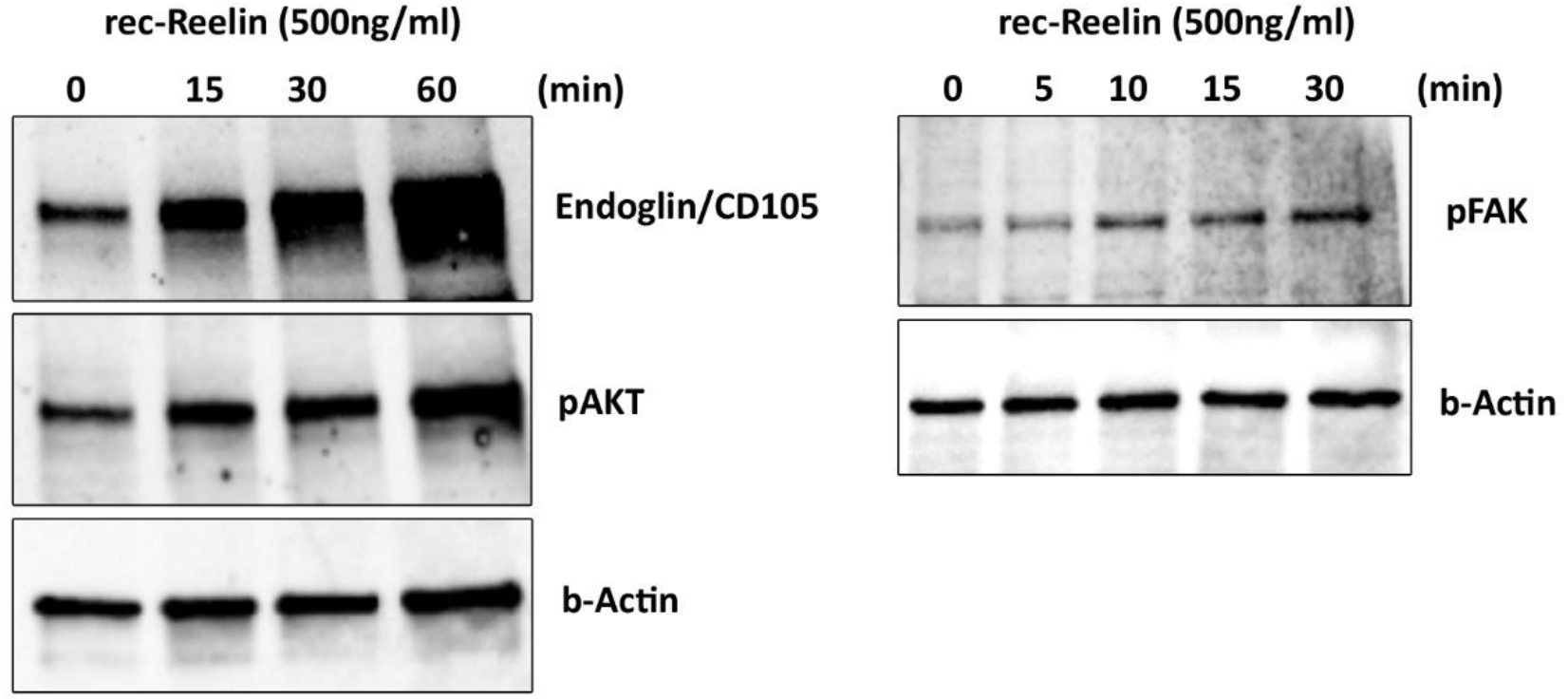
Recombinant Reelin activates focal adhesion and AKT signaling and modulates Endoglin expression in ECs. Representatives immunoblot analyses of ECs stimulated with recombinant Reelin (rec-Reelin, 1 µg/mL) for the indicated times. Left panel: time-dependent phosphorylation of FAK following short-term Reelin stimulation (0–30 min). Right panel: Reelin-induced modulation of Endoglin (CD105) expression and AKT phosphorylation (pAKT) at later time points (0–60 min). β-Actin was used as a loading control in all experiments.

### Reelin induces Endoglin (CD105) expression in ECs

Given the central role of AKT- and focal adhesion–dependent signaling in endothelial activation, vascular remodeling, and cardiovascular disease, we next examined the expression of Endoglin (CD105), a TGF-β co-receptor implicated in angiogenesis and vascular remodeling and frequently associated with early endothelial phenotypic modulation ^39-41^. Reelin stimulation induced a significant upregulation of Endoglin expression in ECs compared with untreated controls (Figure 2A). This response temporally coincided with AKT activation and alterations in FAK-associated signaling, pathways known to regulate endothelial survival, mechanotransduction, and cytoskeletal remodeling within the vascular wall. The coordinated increase in Endoglin expression together with activation of AKT and focal adhesion signaling supports the acquisition of an activated, remodeling-prone endothelial phenotype. Importantly, these changes occurred without evidence of a complete endothelial-to-mesenchymal transition. Rather than indicating full EndMT, our findings are consistent with a state of enhanced endothelial plasticity characterized by selective modulation of remodeling-associated markers. Such partial phenotypic modulation has been described in pathological vascular contexts, where ECs adopt dynamic and reversible changes that contribute to vascular remodeling without undergoing complete mesenchymal conversion. Collectively, these data indicate that Reelin engages signaling pathways relevant to cardiovascular pathophysiology, promoting endothelial activation and plasticity in a manner compatible with vascular remodeling rather than full mesenchymal transition.

### Reelin signaling in ECs is independent of direct VEGFR2 receptor engagement

Given the extensive overlap between Reelin- and VEGF-activated signaling cascades in ECs, both of which are critically implicated in vascular homeostasis and cardiovascular disease, we examined whether Reelin receptors physically associate with VEGFR2 at the plasma membrane. To address this question, co-immunoprecipitation assays were performed using an anti-VEGFR2 antibody, followed by immunoblotting for VEGFR2 and the canonical Reelin receptors ApoER2 and VLDLR.

Immunoblot analysis confirmed efficient and specific immunoprecipitation of VEGFR2, with a strong signal detected in both the input lysate and the VEGFR2 immunoprecipitate, and no signal observed in control immunoprecipitates, indicating minimal nonspecific binding (Figure 3). ApoER2 and VLDLR were readily detectable in input lysates, confirming their expression in ECs; however, neither receptor was detected in VEGFR2 immunoprecipitates. These data indicate that VEGFR2 does not form a stable molecular complex with ApoER2 or VLDLR under basal conditions.

**Figure 3.**
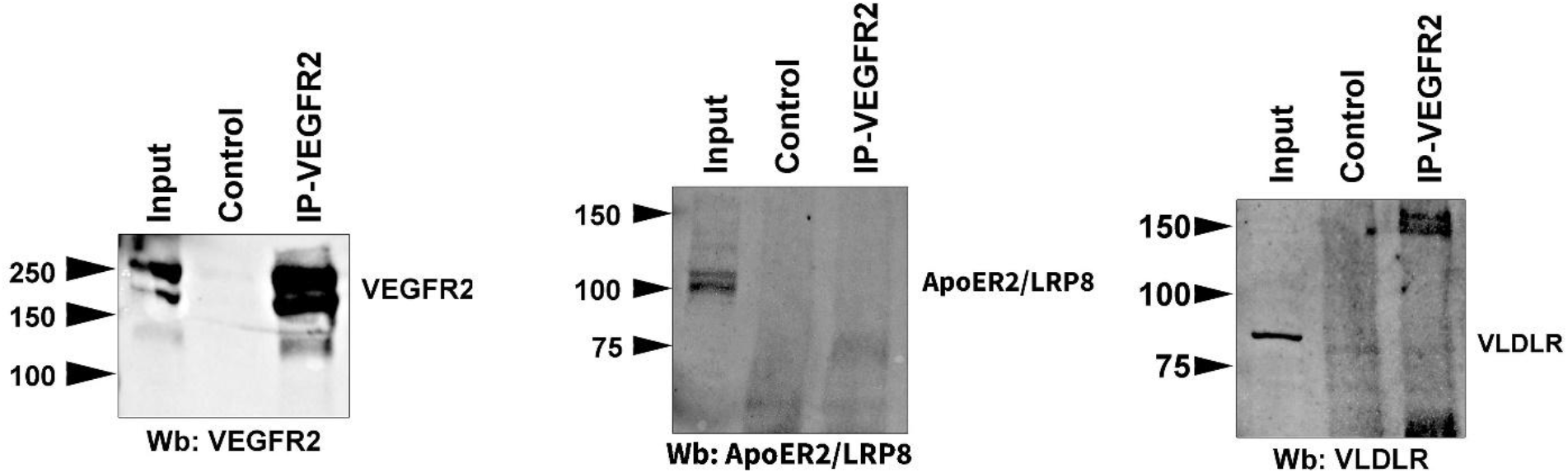
VEGFR2 does not physically associate with Reelin receptors in ECs. EC lysates were subjected to immunoprecipitation (IP) using an anti-VEGFR2 antibody or an isotype-matched control antibody, followed by immunoblot (WB) analysis as indicated. Left panel (WB: VEGFR2): Input lysate shows robust VEGFR2 expression. VEGFR2 is efficiently and specifically immunoprecipitated, while no signal is detected in the control IP, confirming the specificity of the assay. Middle panel (WB: LRP8): LRP8 (ApoER2) is readily detectable in the input lysate, indicating its expression in ECs. No LRP8 signal is observed in the VEGFR2 immunoprecipitate, demonstrating the absence of a physical association between VEGFR2 and LRP8. Right panel (WB: VLDLR): VLDLR is present in the input lysate but is not detected in the VEGFR2 immunoprecipitate, indicating that VEGFR2 does not interact with VLDLR.

Taken together, these findings suggest that Reelin-induced endothelial signaling does not require direct physical interaction with VEGFR2. Instead, Reelin and VEGF pathways are likely to converge downstream of receptor activation, at the level of shared intracellular signaling nodes relevant to cardiovascular biology, including AKT, FAK, and regulators of endothelial cytoskeletal dynamics and cell–cell junctions. This receptor-level independence suggests that Reelin may modulate endothelial plasticity and vascular remodeling through VEGFR2-independent mechanisms that could be particularly relevant in pathological settings such as atherosclerosis, vascular inflammation, and maladaptive angiogenesis

### Reelin alters endothelial migratory behavior in a scratch wound assay

To determine whether Reelin-dependent signaling translates into functionally relevant changes in endothelial behavior with potential implications for vascular pathology, we assessed EC migration using a scratch wound–healing assay. Confluent endothelial monolayers were treated with recombinant Reelin, and wound closure was monitored over time in comparison with untreated controls. Immediately after scratch generation (0 h), control monolayers exhibited a sharply demarcated wound edge, with ECs displaying a typical cobblestone morphology and preserved intercellular junctions (Figure 4). At about 18 h, control cells achieved substantial wound closure, characterized by coordinated advancement of cohesive multicellular sheets and high cell density within the denuded area, consistent with collective migration and/or proliferation-driven repair. In contrast, Reelin-treated ECs exhibited a distinct migratory response. Although cells actively entered the wound area, overall gap closure at 18 h was significantly reduced relative to controls. Reelin-treated ECs adopted an elongated, spindle-shaped morphology and migrated predominantly as single cells or loosely connected groups rather than as organized multicellular fronts. Accordingly, wound margins appeared less compact, and cell density within the scratch area remained lower than in control conditions (Figure 4). These findings indicate that Reelin does not simply enhance endothelial wound repair but instead induces a qualitative shift in migratory mode, favoring individual cell motility over collective movement. Such a shift toward increased endothelial plasticity and reduced intercellular cohesion may be particularly relevant in pathological settings of vascular remodeling, including atherosclerosis, restenosis, and maladaptive angiogenesis, where dysregulated endothelial migration contributes to disease progression.

**Figure 4.**
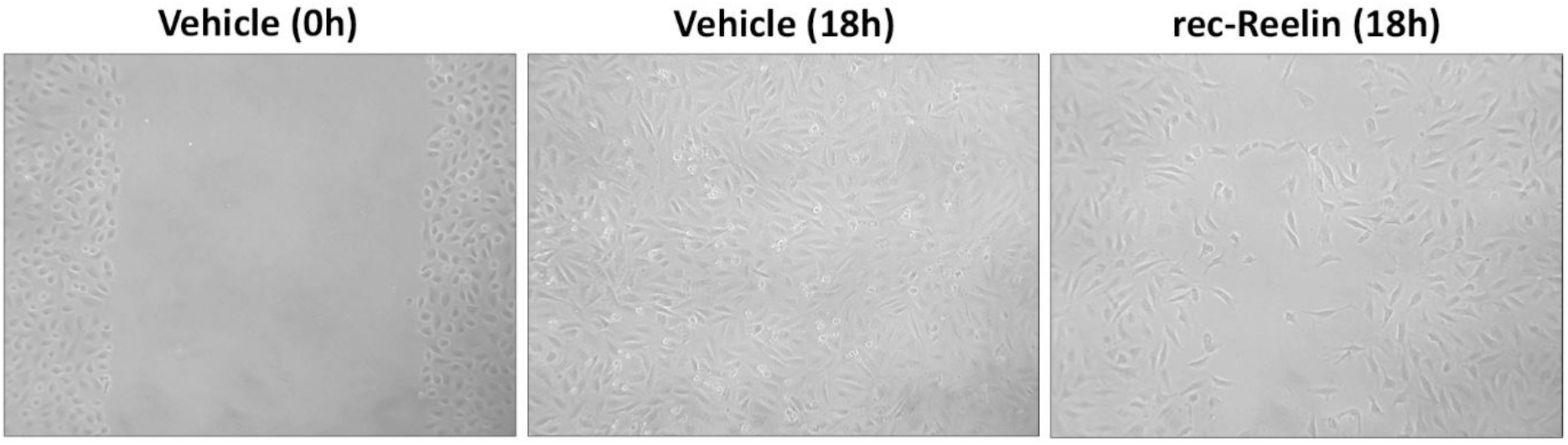
Reelin alters endothelial migratory dynamics in a scratch wound-healing assay. Representative phase-contrast images of EC monolayers subjected to a scratch assay. Images show control cells immediately after scratch formation (Vehicle 0 h) and after 18 h of incubation (Vehicle 18 h), compared with reelin-treated cells after 18 h (rec-Reelin 18 h). Control cells exhibit substantial wound closure characterized by dense, coordinated cell migration into the denuded area. In contrast, reelin-treated cells display reduced gap closure and migrate predominantly as elongated, individually motile cells, resulting in lower cell density within the wound area.

### Reelin silencing disrupts endothelial junctional organization

To further investigate the role of Reelin in endothelial phenotype, we examined the effects of endogenous Reelin depletion in primary ECs. Gene silencing experiments performed in HUVECs revealed measurable alterations in EC morphology and organization, supporting a role for basal Reelin signaling in the maintenance of endothelial homeostasis (data not shown). Notably, the magnitude of these effects showed variability across independent experiments, likely reflecting sensitivity to experimental conditions such as cell confluency and baseline endothelial state. Consistent with these observations, stimulation with recombinant Reelin did not result in statistically significant changes in total CD31/PECAM-1 expression across the analyzed time points, indicating that Reelin does not markedly regulate endothelial junctional protein levels at the protein level under these conditions. Together, these findings suggest that Reelin primarily acts as a context-dependent modulator of endothelial phenotype rather than a direct regulator of junctional protein expression. A schematic representation summarizing the proposed model of Reelin signaling in ECs is shown in Figure 5, highlighting the preferential activation of non-canonical FAK- and AKT-dependent pathways and their association with endothelial plasticity and remodeling.

**Figure 5.**
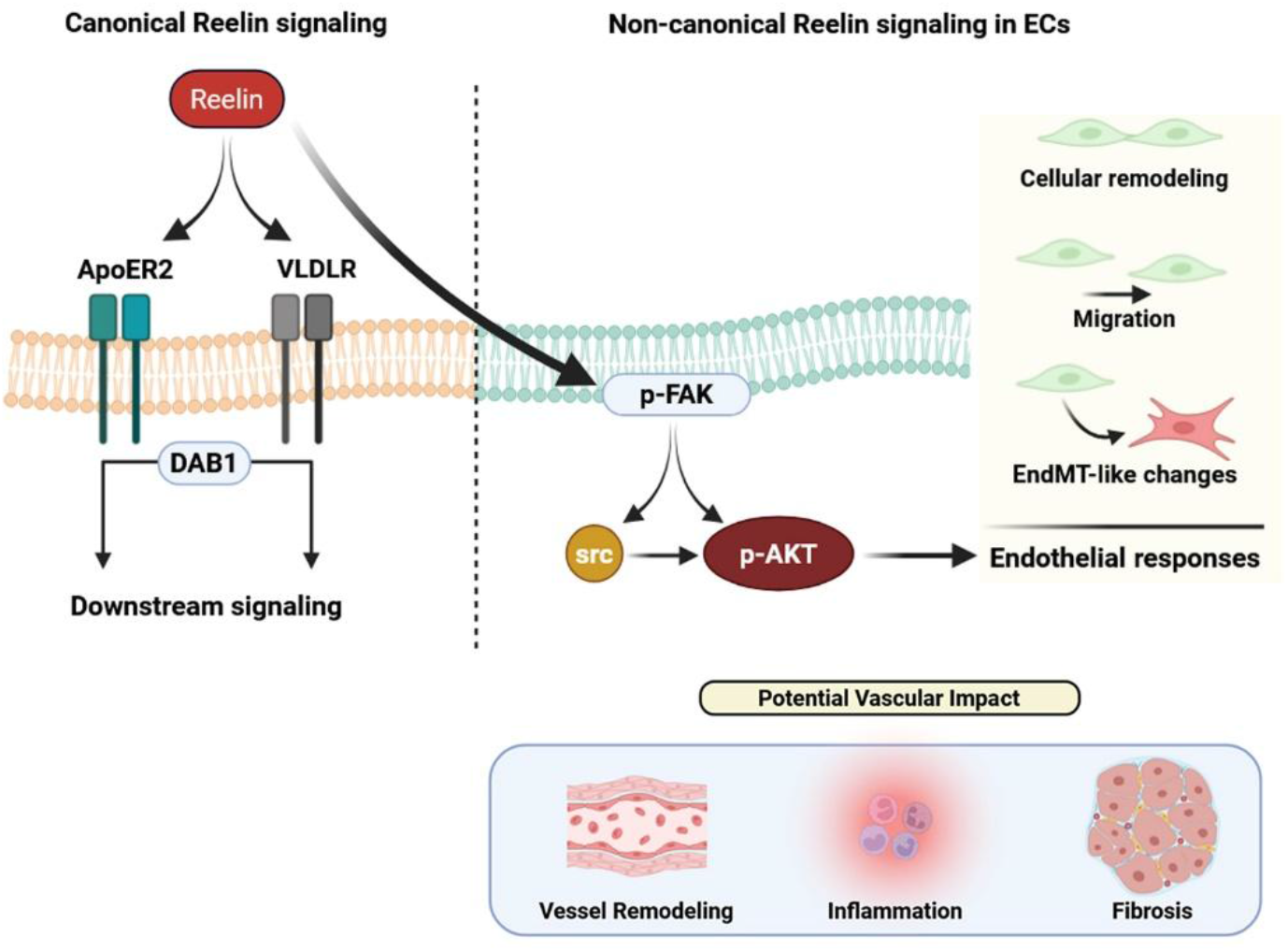
Non-canonical Reelin signaling drives endothelial plasticity and EndMT-like responses. Recombinant Reelin engages its endothelial receptors with limited activation of the canonical DAB1-dependent pathway, resulting in a signaling output that selectively recruits non-canonical effectors. Reelin preferentially triggers a FAK- and AKT-centered signaling cascade, promoting cytoskeletal reorganization, junctional remodeling, and enhanced endothelial migratory capacity. These responses are associated with partial phenotypic modulation consistent with increased endothelial plasticity rather than complete endothelial-to-mesenchymal transition (Created in BioRender).

## Discussion

In the present study, we identify Reelin as a previously unrecognized regulator of endothelial signaling, plasticity, and junctional organization, thereby extending its biological relevance from the nervous system to the cardiovascular endothelium. Our data demonstrate that Reelin modulates endothelial phenotype through non-canonical signaling pathways and promotes partial and dynamically regulated phenotypic modulation, rather than a full mesenchymal transition. Collectively, these findings position Reelin as a novel component of the endothelial regulatory network with potential relevance to vascular remodeling and cardiovascular disease. Endothelial plasticity is increasingly recognized as a fundamental mechanism underlying vascular adaptation and pathology. Partial and reversible EndMT programs, rather than complete mesenchymal conversion, contribute to endothelial dysfunction, inflammation, and fibrotic remodeling in cardiovascular diseases such as atherosclerosis, cardiac fibrosis, pulmonary hypertension, and valvular disease ^19, 42, 43^. In this context, our observation that Reelin induces selective EndMT-associated features without full loss of endothelial identity suggests a modulatory role in endothelial state transitions, potentially enabling adaptive remodeling while predisposing to dysfunction when dysregulated. Importantly, our data do not support complete EndMT conversion but rather indicate a shift toward a remodeling-prone endothelial state. Junctional destabilization is a hallmark of endothelial dysfunction and promotes increased vascular permeability, leukocyte extravasation, and pro-inflammatory vascular remodeling ^44-46^. Our data suggest that Reelin contributes to the maintenance of endothelial barrier integrity, supporting a homeostatic role in preserving endothelial architecture under basal conditions. Despite these functional insights, the precise molecular mechanisms by which Reelin regulates endothelial behavior remain incompletely elucidated. While canonical Reelin signaling pathways involving ApoER2/VLDLR and Dab1 are well characterized in neuronal contexts ^25, 47-49^, our findings indicate that endothelial Reelin signaling exhibits limited canonical engagement with a signaling output predisposed toward non-canonical effectors, particularly FAK and AKT. This mechanistic gap is particularly relevant in the cardiovascular system, where endothelial signaling is tightly modulated by shear stress, inflammatory mediators, and metabolic cues ^2, 50-52^.

A potentially informative aspect of our findings is the apparent lack of direct interaction between the canonical Reelin receptors, and VEGFR2, the principal mediator of VEGF-driven endothelial activation. This suggests that Reelin regulates endothelial behavior through signaling pathways that are mechanistically distinct from classical angiogenic cascades. Rather than modulating VEGF responsiveness, Reelin may act in parallel signaling domains that primarily govern junctional organization, cytoskeletal dynamics, and endothelial cohesion. Such spatial segregation could involve specialized membrane microdomains, such as caveolae, which are known to organize endothelial receptors and signaling molecules into discrete hubs in ECs. Interestingly, previous work in brain ECs has suggested that Reelin can localize to caveolae, supporting the plausibility of compartmentalized signaling in the endothelium ^53^. Further studies will be required to determine whether caveolae or similar structures facilitate compartmentalized Reelin signaling and how this pathway interfaces with canonical angiogenic networks in cardiovascular disease.

From a translational perspective, altered Reelin expression or signaling could contribute to endothelial dysfunction in cardiovascular disease states characterized by chronic inflammation, disturbed flow, or ischemic injury. This hypothesis is supported by recent research demonstrating that circulating Reelin promotes atherosclerosis by enhancing endothelial adhesion molecule expression, leukocyte recruitment, and vascular inflammation, whereas Reelin deficiency attenuates lesion development ^35^. These findings suggest that Reelin is not only a regulator of endothelial plasticity and junctional organization, as demonstrated in this study, but may also influence inflammatory processes that drive vascular remodeling and atherogenesis. The present study extends this concept by indicating that Reelin promotes a remodeling-associated endothelial phenotype characterized by enhanced migratory capacity and focal adhesion signaling, without inducing full mesenchymal transition. In contrast, the therapeutic modulation of Reelin signaling could be considered a strategy for the preservation of endothelial junctional integrity or the restraint of maladaptive endothelial plasticity during vascular remodeling.

## Authors’ contributions

A.D., and D.M.S. designed the experiments; A.D. and M.T.V. secured the funding; D.M.S., and S.N. conducted the experiments and analyzed the data; A.D. wrote the manuscript; A.F. and L.M. provided critical revision of the manuscript.

## Data availability

Raw data are available on request from the corresponding author

